# Regular caffeine intake attenuates REM sleep promotion and sleep quality in healthy men

**DOI:** 10.1101/2020.09.18.291039

**Authors:** Janine Weibel, Yu-Shiuan Lin, Hans-Peter Landolt, Christian Berthomier, Marie Brandewinder, Joshua Kistler, Sophia Rehm, Katharina M. Rentsch, Martin Meyer, Stefan Borgwardt, Christian Cajochen, Carolin F. Reichert

**Author notes:** Authors contributed equally to this work (Christian Cajochen, Carolin F. Reichert). Address for correspondence: Christian Cajochen, PhD, Centre for Chronobiology, Psychiatric Hospital of the University of Basel, Wilhelm Klein-Strasse 27, CH-4002 Basel.

## Abstract

Acute caffeine intake can attenuate homeostatic sleep pressure and worsen sleep quality. Besides, caffeine intake – particularly in high doses and close to bedtime – may also affect circadian-regulated REM sleep promotion, an important determinant of subjective sleep quality. However, it is not known whether such changes persist under chronic caffeine consumption during daytime. Twenty male caffeine consumers (26.4 ± 4 years old, habitual caffeine intake 478.1 ± 102.8 mg/day) participated in a double-blind crossover study. Each volunteer completed a caffeine (3 × 150 mg caffeine daily), a withdrawal (3 × 150 mg caffeine for eight days then placebo), and a placebo condition. After ten days of controlled intake and a fixed sleep-wake cycle, we recorded 8 h of electroencephalography starting 5 h after habitual bedtime (i.e., start on average at 04:22 am which is around the peak of circadian REM sleep promotion). A 60 min evening nap preceded each sleep episode and reduced high sleep pressure levels. While total sleep time and sleep architecture did not significantly differ between the three conditions, REM latency was longer after daily caffeine intake compared to both placebo and withdrawal. Moreover, the accumulation of REM sleep proportion was slower, and volunteers reported more difficulties at awakening after sleep and feeling more tired upon wake-up in the caffeine condition compared to placebo. Our data indicate that besides acute also regular daytime caffeine intake affects REM sleep regulation in men. We have evidence that regular caffeine intake during daytime weakens circadian sleep promotion when compared to placebo. Moreover, the observed caffeine-induced deterioration in the quality of awakening may suggest a potential motive to reinstate caffeine intake after sleep.

## 1. Introduction

Caffeine has been used for centuries (Camandola et al., 2019) and is considered to be today’s most popular stimulating substance around the globe (Fredholm et al., 1999). Approximately 80% of the world’s population consume caffeinated aliments day by day (Heckman et al., 2010). The common pattern of caffeine intake in the morning and afternoon (Martyn et al., 2018; Lieberman et al., 2019) most likely originates from the motive to achieve both benefits in daytime alertness (Smith, 2002; Einöther and Giesbrecht, 2013) and a good subjective nighttime sleep quality despite of previous stimulant consumption (Snel and Lorist, 2011). These double-edged alerting but sleep-disrupting effects of caffeine have been traced back to its impact on the homeostatic component of sleep-wake regulation (Landolt, 2008). By antagonizing adenosine (Fredholm et al., 1999), caffeine dampens homeostatic sleep need (Landolt, 2008), as evident in caffeine-induced reductions of waking EEG theta activity (Landolt et al., 2004), slow-wave sleep (SWS), and slow-wave activity (SWA), while it increases activity in the sigma range (Landolt et al., 1995a). Importantly, structure and intensity of sleep are not only determined by homeostatic sleep need but also by the circadian timing system (Borbély, 1982; Lazar et al., 2015). One of the most prominent circadian sleep features is REM sleep propensity, which usually peaks in the morning around 2 h after the nadir of core body temperature (Dijk and Czeisler, 1995). This natural peak right before usual wake time may facilitate the re-arousal of the brain from sleep (Dijk and Czeisler, 1995) and may contribute to REM-sleep’s substantial promotion of good subjective sleep quality (Akerstedt et al., 1994; Della Monica et al., 2018). Since the choice to consume caffeine might also strongly depend on an individual’s sleep quality, it remains to be established how daily caffeine intake impacts on REM sleep and its role in promoting sleep quality.

There is evidence for acute effects of caffeine on REM sleep. In animal models the perfusion of the *sleep factor* adenosine increased time spent in REM sleep (Portas et al., 1997; Basheer et al., 1999). Regarding nighttime sleep in humans, some studies – particularly those utilizing high caffeine dosages (but see (Bonnet and Arand, 1992; Drake et al., 2013)) – report a caffeine-induced reduction of REM sleep duration (Brezinova, 1974; Nicholson and Stone, 1980; Robillard et al., 2015) or shift in REM sleep episodes (Karacan et al., 1976; Nicholson and Stone, 1980) while others did not report any differences in REM sleep after caffeine administration (Bonnet and Arand, 1992; Landolt et al., 1995a; Landolt et al., 1995b; Drapeau et al., 2006; Drake et al., 2013). When sleep was initiated around the peak of REM sleep propensity (i.e., 1 h after habitual wake time) following one night of sleep loss, caffeine intake right before bedtime reduced REM sleep at the cost of wakefulness (Carrier et al., 2007; Carrier et al., 2009). Whether such effects during daytime recovery sleep can also be observed under conditions of regular daily daytime caffeine intake remains unknown.

Regular daytime caffeine intake can lead to adaptations and thus mitigate both caffeine-induced wake-promotion and nighttime sleep-disturbances. Such adaptations are represented in the occurrence of withdrawal symptoms within around 36 h after acute cessation of caffeine, and comprise increased sleepiness (James, 1998; Weibel et al., 2020b), enhanced waking EEG theta activity (Sigmon et al., 2009), and reduced sleep EEG sigma activity (Weibel et al., 2020a). Thus, comparing withdrawal-induced effects on sleep against long-term abstinence and habitual use in the same individuals represents a valid tool to reliably estimate the consequences of daily caffeine intake. For this report, we took advantage of an existing data set with exactly these conditions, in which the start of each of the sleep episodes was individually scheduled around the circadian peak of REM sleep promotion. Previous sleep-wake history, light input, posture, meal intake, and circadian phase (estimated by dim-light melatonin onset) were carefully controlled and did not differ between conditions or within individuals (Weibel et al., 2020b). Based on the evidence summarized above, we explored whether the duration and timing of REM sleep change in response to daily daytime caffeine intake (over 10 days) and acute caffeine withdrawal, compared to a long-term (10-day) placebo baseline. In a second step, we tested whether changes in REM sleep relate to differences in subjective sleep quality.

## 2. Methods

### 2.1. Volunteers

Data sets were available from a total of 20 male study volunteers between 18 and 32 years old. All participants were regular caffeine consumers with a daily intake between 300 and 600 mg assessed by a survey tool based on (Bühler et al., 2013) and its caffeine content classified according to (Snel and Lorist, 2011). Prior to study participation, all volunteers were screened for good health assessed by self-report questionnaires and a medical check performed by a study physician. Individuals reporting a BMI < 18 or > 26 were excluded. Good sleep quality was ensured by the Pittsburgh Sleep Quality Index (PSQI; score ≤ 5) (Buysse et al., 1989) and a polysomnography (PSG) during which we screened for sleep apnea (index > 10), period leg movements (index > 15/h), and poor sleep efficiency (SE < 70%). Smoking, drug use or extreme chronotype (score ≤ 30 or ≥ 70 in the Morningness-Eveningness Questionnaire (Horne and Ostberg, 1976)) resulted in the exclusion of participants. In addition, volunteers were not allowed to engage in shift work (< three months prior to study) or to travel across more than two time zones (< one month prior to study). In order to minimize the potential confounding by the menstrual cycle and the use of oral contraceptives on sleep (Shechter and Boivin, 2010) and caffeine elimination (Abernethy and Todd, 1985; Balogh et al., 1995), female individuals were not included in the present study. Demographical data of the study sample can be found in Table 1.

**Table 1.** Demographical data of the study sample.

| Sample characteristics (N=20) | Mean ± SD |
| --- | --- |
| Years of age | 26.4 ± 4.0 |
| Habitual caffeine intake (mg/day) | 478.1 ± 102.8 |
| BMI (kg/m<sup>2</sup>) | 22.7 ± 1.4 |
| Chronotype (MEQ) | 52.8 ± 8.7 |
| Sleep quality (PSQI) | 2.8 ± 1.4 |
| Habitual bedtime (hh:mm)<sup>1</sup> | 23:21 ± 00:49 |
| Habitual sleep duration (hh:mm)<sup>1</sup> | 07:28 ± 00:25 |

Notes. BMI: Body Mass Index; MEQ: Morningness-Eveningness Questionnaire (Horne and Ostberg, 1976); PSQI: Pittsburgh Sleep Quality Index (Buysse et al., 1989); <sup>1</sup>self-reported.

### 2.2. Design and Protocol

We employed a double-blind crossover study with three treatments: a caffeine, a withdrawal, and a placebo condition. Random permutations were performed to assign the volunteers to the order of the three conditions, for more details see (Weibel et al., 2020b). As depicted in Fig. 1a, each condition comprised an ambulatory part of nine days which was followed by an in-lab part of 43 h. In each condition, volunteers swallowed identical appearing gelatin capsules three times daily (45 min, 255 min, and 475 min after awakening) containing either caffeine (150 mg; Hänseler AG, Herisau, Switzerland) or placebo (mannitol; Hänseler AG, Herisau, Switzerland). To induce caffeine withdrawal in the withdrawal condition, the first capsule on day nine of treatment contained caffeine but was followed by placebo capsules for the remaining administrations. The dosage and timing of administrations was chosen based on previous studies which investigated tolerance development to the effects of caffeine and its cessation (James, 1998; Keane and James, 2008) and to represent an everyday situation of most coffee consumers (Martyn et al., 2018; Lieberman et al., 2019). Volunteers were instructed to abstain from all caffeine sources during the entire study duration. Compliance to the aforementioned regimen was verified by assessing caffeine levels in fingertip sweat collected prior to habitual bedtime. Moreover, volunteers were instructed to keep a regular sleep-wake rhythm (± 30 min of self-selected bedtime and waketime, 8 h in bed) and to avoid naps. Their compliance was monitored by wrist actimetry (Actiwatch, Cambridge Neurotechnology, Cambridge, United Kingdom) with concurrent sleep logs. Sleep schedule remained constant across all three conditions except for three volunteers (two volunteers: +30 min in caffeine compared to placebo and withdrawal conditions; one volunteer: −30 min in placebo compared to caffeine and withdrawal conditions).

**Fig. 1.**
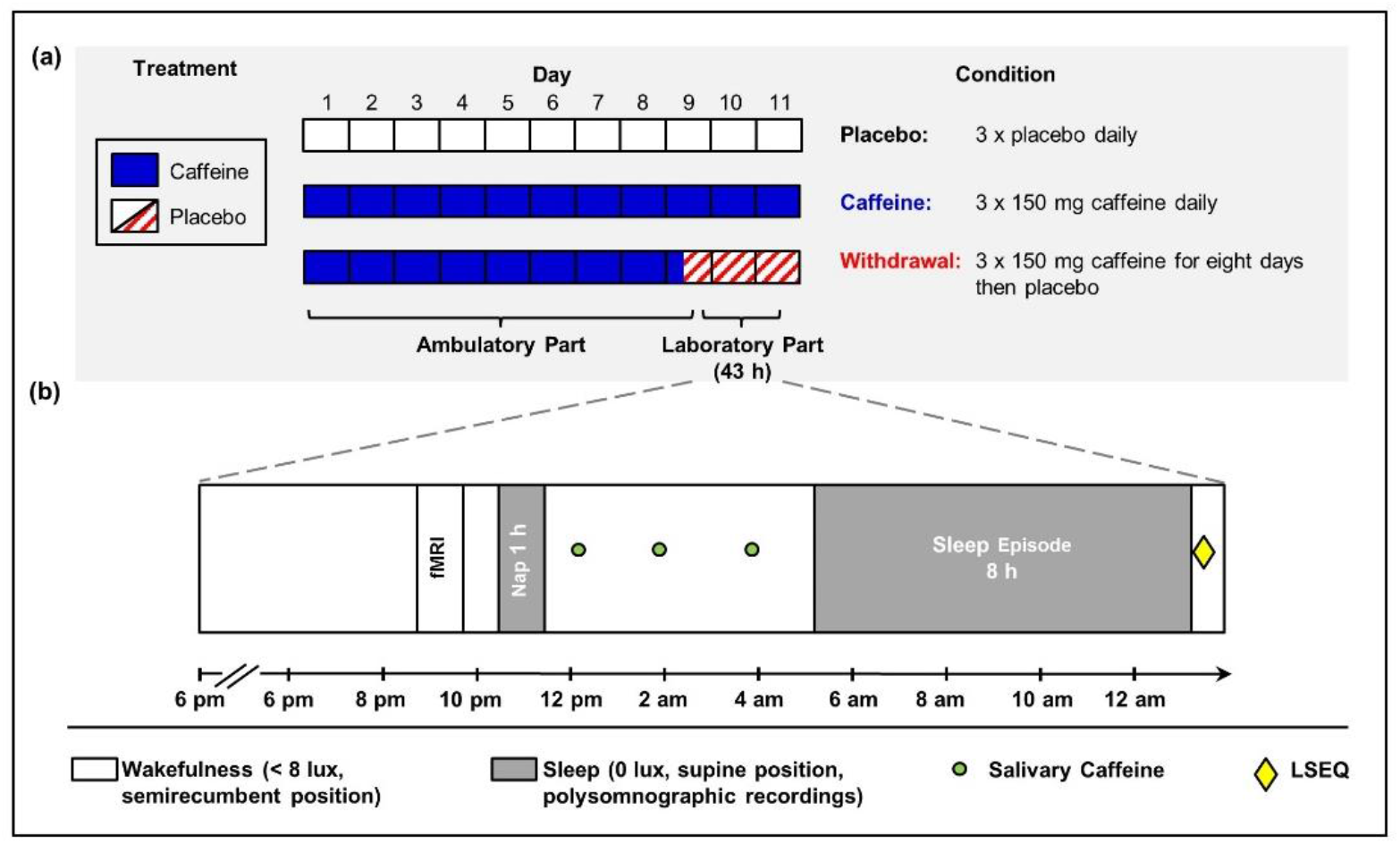
Illustration of the research protocol (adapted from (Weibel et al., 2020b)). **(a)** Each participant took part in a placebo, a caffeine, and a withdrawal condition consisting of an ambulatory part of nine days and an in-lab part of 43 h. **(b)** The in-lab protocol started with a baseline night scheduled to volunteers’ habitual bedtime. On the following day, we scheduled a 1-hour nap in the evening and salivary caffeine levels were collected in regular intervals. Five hours after usual bedtime, an 8-hour sleep episode was scheduled, and subjective sleep quality was assessed afterwards.

As illustrated in Fig. 1b, on day nine of treatment, volunteers reported to the laboratory 5.5 h prior to habitual bedtime. Upon arrival, PSG electrodes were fitted and a sleep episode of 8 h was scheduled at the volunteer’s habitual bedtime. The following day, saliva was regularly collected in order to quantify the levels of caffeine and its main metabolites. In the evening, we scheduled a one-hour nap (for details see (Weibel et al., 2020b)), which was followed by another wakefulness episode. To minimize masking effects during the in-lab protocol, no time-of-day information was provided, communication was restricted to staff members, light (scheduled wakefulness: < 8 lux; sleep episode: 0 lux), and posture (scheduled wakefulness: semirecumbent; sleep episode: supine) was controlled.

Approximately 5 h after volunteers’ habitual bedtime (mean: 04:22 am; range: 03:15 – 05:15 am), a sleep episode of 8 h was scheduled, which corresponds to the expected circadian peak of REM sleep promotion (Dijk and Czeisler, 1995; Münch et al., 2005), and was 13.5 and 44.5 h after the last caffeine intake in the caffeine and withdrawal condition, respectively. Upon awakening, volunteers rated their subjective sleep quality by the Leeds Sleep Evaluation Questionnaire (LSEQ) (Parrott and Hindmarch, 1978).

Approval was obtained from the local ethics committee (Ethikkommission Nordwest- und Zentralschweiz) and the study was conducted in accordance with the declaration of Helsinki. All volunteers provided a written informed consent and received financial compensation for study participation.

### 2.3. Caffeine Levels

Salivary caffeine levels were repeatedly assessed in intervals of approximately two hours throughout the in-lab protocol to check compliance with the treatment requirements. Here we focus on the levels within 5 h prior to the sleep episode. After saliva collection, samples were immediately stored at 5°C, later centrifuged at 3000 rpm for 10 min, and subsequently frozen at – 24°C until assayed by liquid chromatography coupled to tandem mass spectrometry. The data of one volunteer collected in the withdrawal condition was lost.

### 2.4. Subjective Sleep Quality

To assess subjective sleep quality, volunteers completed the LSEQ questionnaire (Parrott and Hindmarch, 1978) right after the end of the 8-h nighttime sleep episode. Volunteers rated 10 items on visual analogue scales consisting of four different scales, i.e. getting to sleep (GTS), quality of sleep (QOS), awake following sleep (AFS), and behavior following wakening (BFW).

### 2.5. Electroencephalographic Recordings

We utilized electroencephalographic recordings to assess sleep structure. Six EEG derivations (F3, F4, C3, C4, O1, O2), two electrooculographic, two electromyographic, and two electrocardiographic electrodes were placed according to the international 10-20 system and referenced online against the linked mastoids (A1, A2). The EEG signal was recorded using V-Amp devices (Brain Products, Gilching, Germany) with a sampling rate of 500 Hz and a filter applied online at 50 Hz.

Sleep staging was performed with an automatic scoring algorithm (ASEEGA, version 4.4.23, PHYSIP, Paris, France) which has been successfully used in previous studies (Reichert et al., 2017; Gaggioni et al., 2019) and has been shown to reach good agreement with manual sleep scoring (Berthomier et al., 2020). The concordance of a subset of manually scored nights (50 nights of the present study sample) with the automatic sleep scoring reached > 80%. The derivation (C4O2) was used for sleep autoscoring. Sleep latencies to specific stages were defined based on the first epoch scored in the respective sleep stage. While we focus in the present paper on REM sleep, we report all other sleep stages as well. All sleep stages are expressed as percentage of total sleep time (TST). Spectral analyses were performed by employing a fast Fourier transform with a Hanning window on consecutive epochs of 30 seconds. Artefacts were automatically rejected, and power spectra in the delta band (0.1-4 Hz) during NREM is reported as a measure for sleep pressure.

Three datasets were excluded from analyses of all-night parameters concerning the entire sleep episode due to technical issues (placebo condition: *n* = 2; caffeine condition: *n* = 1) and one volunteer was excluded from all analyses of REM parameters (placebo condition) based on potential missed REM episode by the algorithm and disagreement with visual scorers.

### 2.6. Statistical Analyses

Data analyses were performed using SAS (version 9.4, SAS Institute, Cary, United States). Values exceeding three times the interquartile range (IQR) from the first and third quartile were treated as extreme outliers and removed from subsequent analyses when attributed to errors in data collection or subsequent handling (REM latency = 1). We applied mixed model analyses of variance with the factors condition (placebo, caffeine, and withdrawal) and time (levels differ per variable). The degrees of freedom were based on the approximation of Kenward and Roger reported in (Kenward and Roger, 1997). Contrasts were calculated by applying the LSMEANS statement and were adjusted for multiple comparisons based on Tukey Kramer. To investigate whether the ratings of sleep quality can be explained by REM sleep parameters, we performed general linear models with the statistics software SPSS (version 24, IBM Corporation, Armonk, NY, USA) including REM sleep parameters as covariates and subject as a random factor. A p-value < 0.05 was used to determine statistical significance. However, the thresholds for the sleep stages and subjective sleep quality were adjusted according to the Bonferroni method (to *p* < 0.0055 and *p* < 0.0125 respectively) due to multiple testing. One volunteer was excluded from all analyses due to noncompliance with the treatment requirements (caffeine condition: *n* = 1).

## 3. Results

### 3.1. Salivary Caffeine Levels

The analyses of caffeine levels assessed within 5 h prior to the sleep episode revealed a significant main effect of condition (*F*_2,55.9_ = 21.79; *p* < 0.0001). Post-hoc comparisons indicated that caffeine levels were still elevated in the caffeine compared to the placebo and withdrawal conditions (*p*_all_ < 0.0001), as presented in Fig. 2. Caffeine levels in the withdrawal condition did not significantly differ from placebo (*p* = 0.984).

**Fig. 2.**
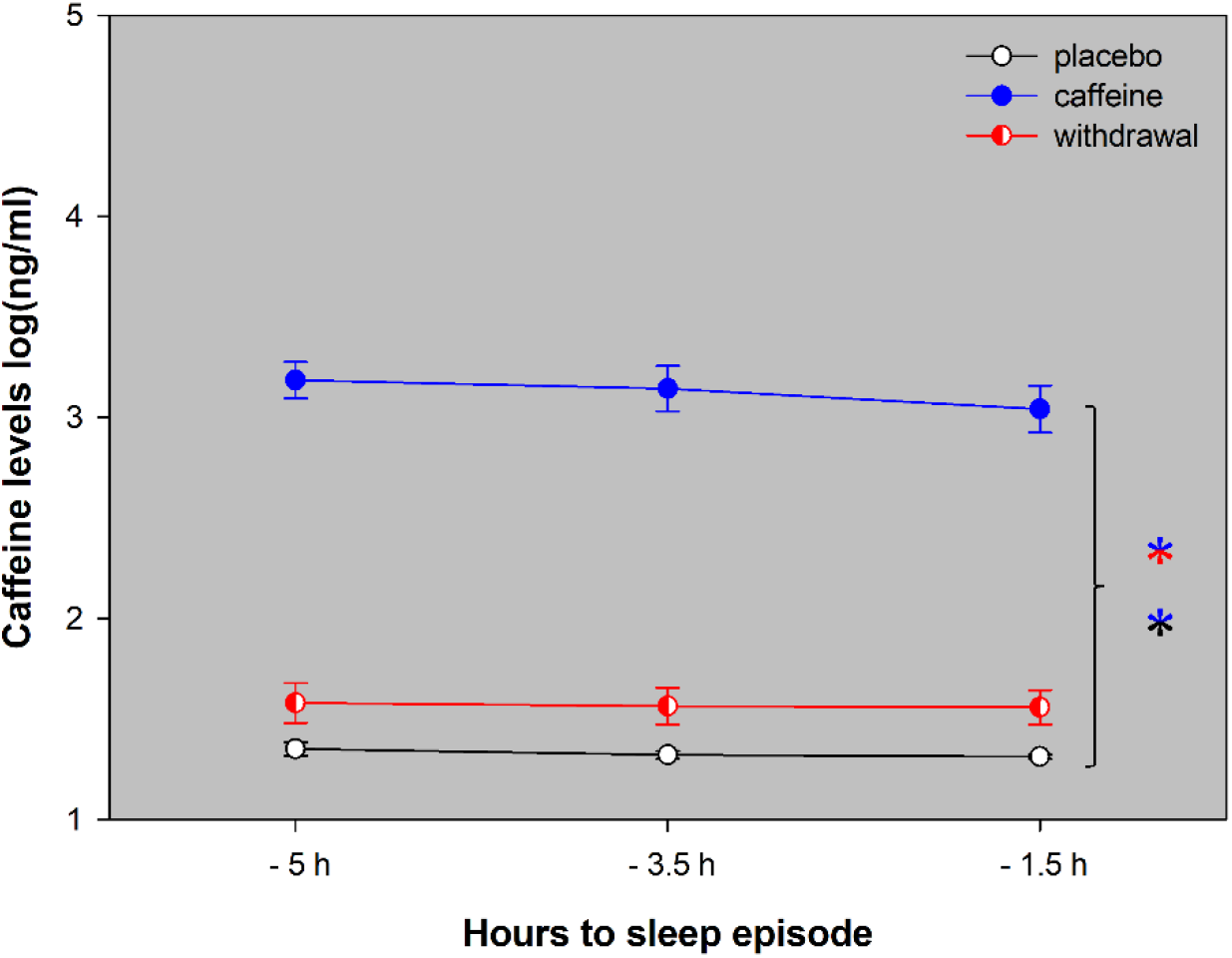
Depicted are the salivary caffeine levels collected within 5 h prior to the sleep episode in the placebo (black open circles), caffeine (blue filled circles), and withdrawal (red semi-filled circles) conditions (means ± standard errors). Overall, caffeine levels were still increased in the caffeine condition compared to both placebo and withdrawal (*p* < 0.05).

### 3.2. Sleep

#### 3.2.1. Electroencephalographic Recordings

We analyzed total sleep time (TST), sleep latencies, and sleep architecture of the 8 h sleep episode, as summarized in Table 2. TST and the proportion of each sleep stage did not significantly differ among the three conditions (*p*_all_ > 0.1). However, in the caffeine condition it took participants longer to enter REM sleep compared to the placebo and withdrawal conditions (*p_all_* < 0.05).

**Table 2.** Sleep variables and results of the electroencephalographic variables.

| Variable | Placebo | Caffeine | Withdrawal | Condition |
| --- | --- | --- | --- | --- |
| TST (min) | 366.19 ± 16.71 | 393.89 ± 13.94 | 393.20 ± 11.23 | $F_{2,35.2} = 2.32, p = 0.113$ |
| SE (%) | 76.79 ± 3.44 | 82.36 ± 2.79 | 82.46 ± 2.30 | $F_{2,35.3} = 2.27, p = 0.118$ |
| N1 (% of TST) | 3.78 ± 0.52 | 4.49 ± 0.79 | 4.20 ± 0.48 | $F_{2,35.9} = 0.40, p = 0.676$ |
| N2 (% of TST) | 44.93 ± 1.80 | 45.57 ± 1.39 | 44.69 ± 1.45 | $F_{2,34.6} = 0.06, p = 0.942$ |
| N3 (% of TST) | 24.21 ± 1.25 | 25.22 ± 1.25 | 24.40 ± 1.31 | $F_{2,35.4} = 0.27, p = 0.763$ |
| REM (% of TST) | 27.82 ± 1.31 | 24.73 ± 1.55 | 26.72 ± 1.11 | $F_{2,34.9} = 1.87, p = 0.169$ |
| SL2 | 11.13 ± 2.29 | 9.13 ± 0.89 | 8.13 ± 1.11 | $F_{2,37.3} = 1.62, p = 0.212$ |
| RL | 53.63 ± 5.53 | 78.74 ± 10.21* | 53.95 ± 6.42 | $F_{2,36.4} = 6.30, p = 0.005$ |
| NA | 8.61 ± 1.30 | 9.22 ± 1.18 | 8.50 ± 1.27 | $F_{2,35.4} = 0.14, p = 0.871$ |

Depicted are the means and standard errors per condition. TST: total sleep time (sum of N1, N2, N3, and REM); SE: sleep efficiency (TST/time in bed); N1: stage 1; N2: stage 2; N3: slow-wave sleep; REM: rapid eye movement sleep; SL2: time from lights-off to first epoch of N2; RL: time from lights-off to first epoch of REM sleep; NA: number of awakenings.
\*significant post-hoc comparisons ( $p < 0.0055$ ; threshold adjusted according to Bonferroni) compared to placebo and withdrawal conditions.

Based on the reductions and shifts of REM sleep reported in previous studies (Karacan et al., 1976; Nicholson and Stone, 1980; Carrier et al., 2007; Robillard et al., 2015), we investigated in a next step whether the accumulation of REM sleep proportion across the sleep opportunity differs among the three conditions. As shown in Fig. 3, on average REM proportion was reduced in the caffeine compared to the placebo condition (main effect of condition: *F*_2,39.1_ = 6.75, *p* = 0.003). This effect was modulated by time (interaction of condition × time: *F*_14,195_ = 1.77, *p* = 0.046) indicating that this caffeine-induced reduction in the caffeine condition was particularly present between 1 to 3 and 4 to 7 h into the sleep opportunity. Importantly, the analyses of the accumulation of wakefulness and SWS (see Fig. 3) did neither yield a significant main effect of condition (wakefulness: *F*_2,36.4_ = 0.49, *p* = 0.615; SWS: *F*_2,38.6_ = 0.94, *p* = 0.401) nor a significant interaction of the factors condition and time (wakefulness: *F*_14,205_ = 0.86, *p* = 0.603; SWS: *F*_14,196_ = 0.54, *p* = 0.908).

**Fig. 3.**
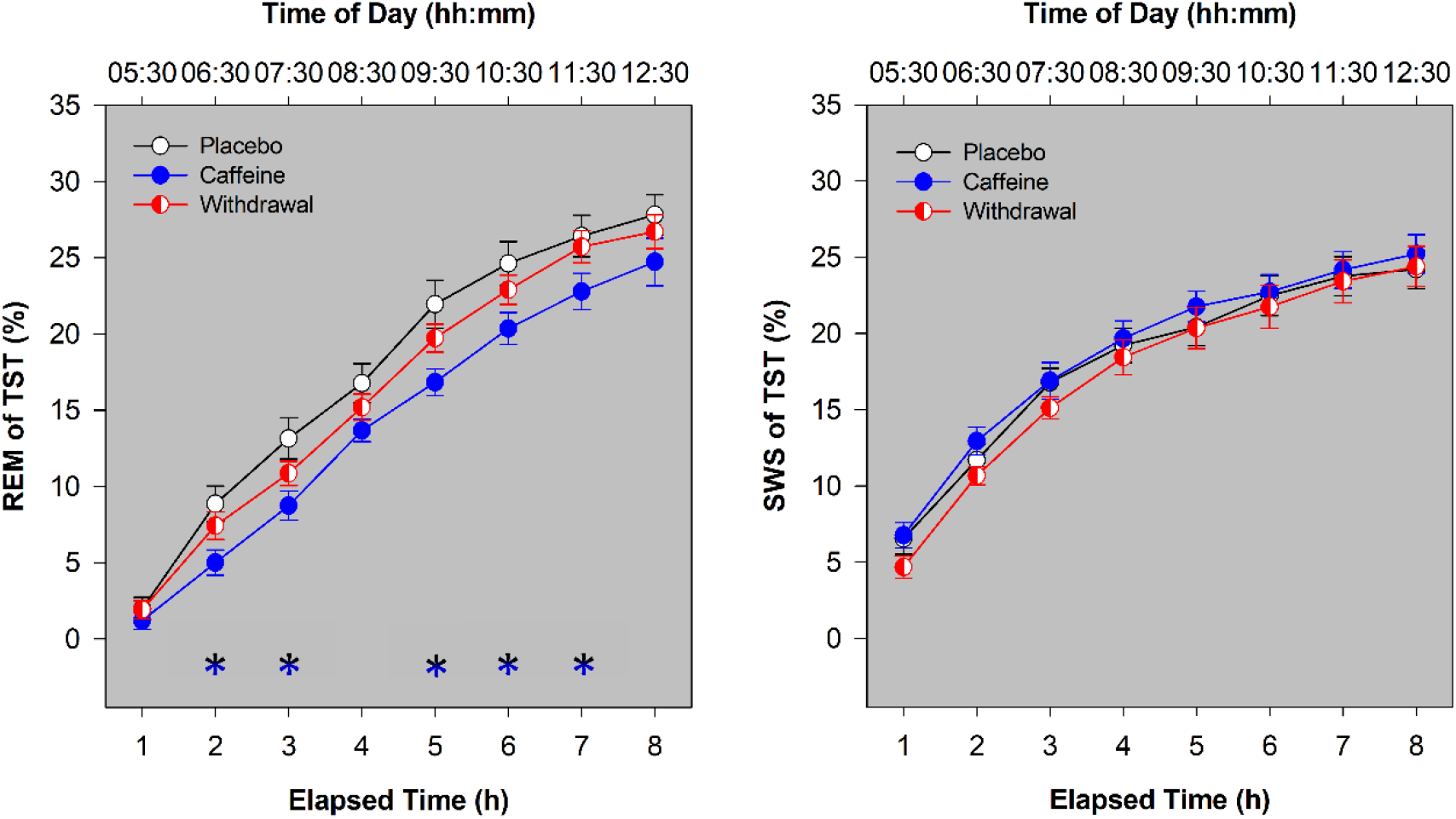
Accumulation of REM and SWS proportion across the sleep opportunity of 8 h. REM (of TST) and SWS (of TST) were collapsed into bins of one hour and accumulated across the sleep episode. Depicted are means and standard errors of the placebo (black open circles), caffeine (blue filled circles), and withdrawal conditions (red semi-filled circles). The color-coded asterisks represent significant (*p*_all_ < 0.05) differences between the placebo and caffeine conditions corrected for multiple comparisons according to (Curran-Everett, 2000).

Lastly, we analyzed the spectral power in the delta band (0.1-4 Hz) across the first three NREM cycles as an indicator for sleep pressure. There were no significant differences between the three conditions in NREM delta power (main effect of condition: *F*_2,26.8_ = 1.19; *p* > 0.3) nor was it modulated by the NREM cycles (interaction condition × time: *F*_4,45.1_ = 0.99; *p* > 0.4).

#### 3.2.2. Subjective Sleep Quality

The parameters and results of subjective sleep quality are presented in Table 3. While the domains *getting to sleep* and *quality of sleep* did not significantly differ between the three conditions (*p*_all_ > 0.05), volunteers reported more difficulty in awakening following sleep (AFS) and feeling more tired following wakening (BFW) in the caffeine compared to placebo condition (*p*_all_ < 0.01). Neither REM sleep duration nor REM (% of TST) had a significant effect on the ratings of AFS (*p*_all_ > 0.4) and BFW (*p*_all_ > 0.3).

**Table 3.**
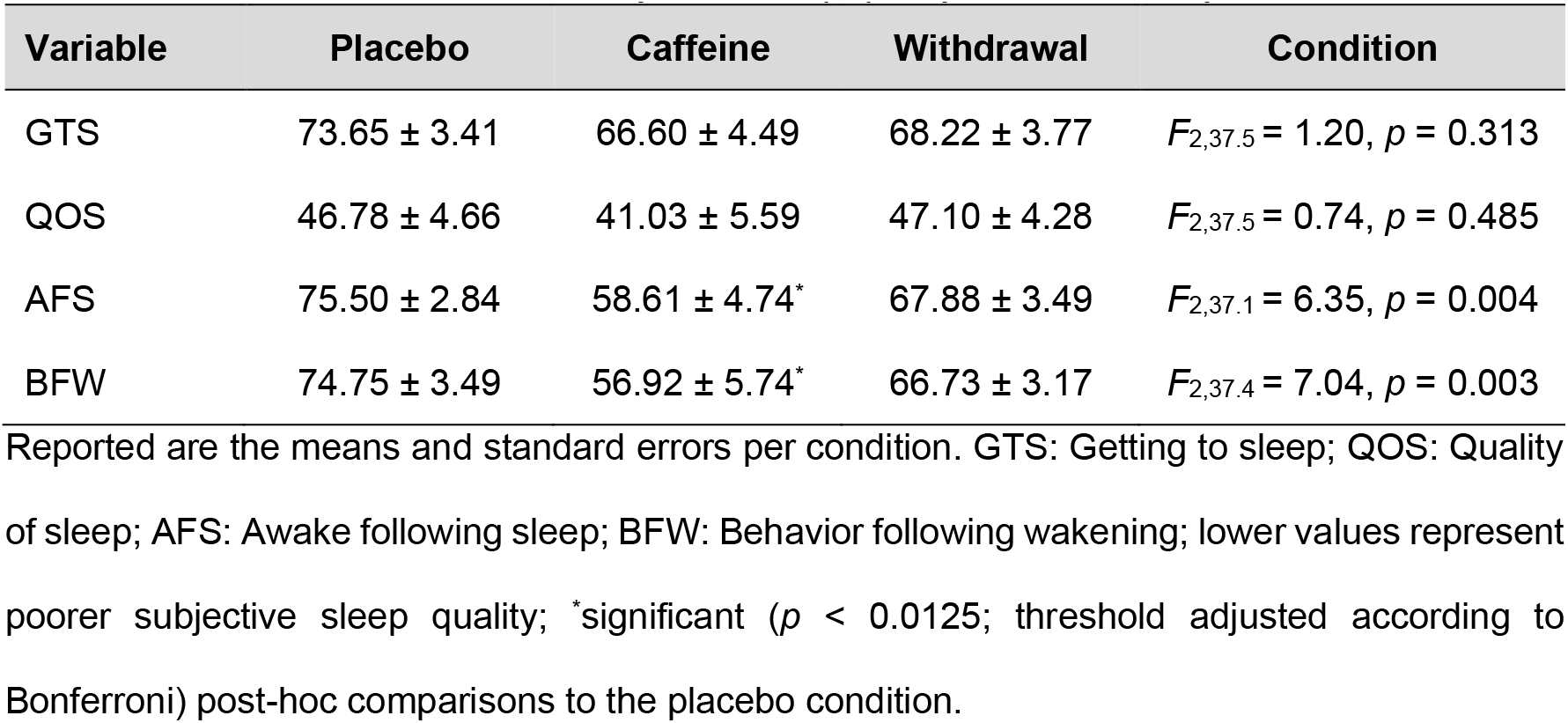
Parameters and results of subjective sleep quality as assessed by the LSEQ.

| Variable | Placebo | Caffeine | Withdrawal | Condition |
| --- | --- | --- | --- | --- |
| GTS | 73.65 ± 3.41 | 66.60 ± 4.49 | 68.22 ± 3.77 | $F_{2,37.5} = 1.20$, $p = 0.313$ |
| QOS | 46.78 ± 4.66 | 41.03 ± 5.59 | 47.10 ± 4.28 | $F_{2,37.5} = 0.74$, $p = 0.485$ |
| AFS | 75.50 ± 2.84 | 58.61 ± 4.74* | 67.88 ± 3.49 | $F_{2,37.1} = 6.35$, $p = 0.004$ |
| BFW | 74.75 ± 3.49 | 56.92 ± 5.74* | 66.73 ± 3.17 | $F_{2,37.4} = 7.04$, $p = 0.003$ |

Reported are the means and standard errors per condition. GTS: Getting to sleep; QOS: Quality of sleep; AFS: Awake following sleep; BFW: Behavior following wakening; lower values represent poorer subjective sleep quality; \*significant ( $p < 0.0125$ ; threshold adjusted according to Bonferroni) post-hoc comparisons to the placebo condition.

## 4. Discussion

To the best of our knowledge, we explored for the first time the effects of regular daytime caffeine intake on REM sleep promotion and subjective quality of sleep. An 8-hour sleep window scheduled in the early morning hours allowed to investigate REM sleep expression at its circadian maximum (Dijk and Landolt, 2019) in a highly controlled laboratory setting. Indicating a reduced circadian promotion of sleep (Dijk and Landolt, 2019), volunteers in the caffeine condition entered REM sleep later compared to both placebo and withdrawal conditions and the accumulation of REM sleep across the night was slower compared to placebo. Thus, the earlier reported REM sleep reductions after caffeine right before bedtime (Carrier et al., 2007; Carrier et al., 2009; Robillard et al., 2015) seem to persist, even if caffeine intake is restricted to daytime and the last intake is 13.5 h apart from lights-off. Moreover, volunteers reported more difficulties in awakening from sleep and feeling more tired after daily caffeine intake compared to continuous placebo intake. Such subtle changes in subjective sleep quality may promote the maintenance of regular caffeine intake under conditions of delayed or shifted sleep.

While daily caffeine intake is highly popular, the effects of regular daytime consumption on sleep-wake regulation are not well understood. One reason might be difficulties to standardize the history of prior caffeine consumption and to control its daily intake. Moreover, a potential adaptation to the continuous availability of caffeine (Bonnet and Arand, 1992; Weibel et al., 2020b), requires a design sophisticated enough to disentangle an inherent insensitivity from a habituation to the stimulant’s effect (James, 1998). In the present report, for which we took advantage from an existing highly controlled laboratory data set, we suggest that regular daytime caffeine intake has the potential to alter REM sleep, indexed as delayed onset and reduced accumulation. As a few studies suggest an adaptation of several nighttime sleep features (Bonnet and Arand, 1992) and of the circadian timing of melatonin onset (Weibel et al., 2020b) in response to regular caffeine intake, the present results indicate that the habituation of REM sleep promotion to caffeine might either follow another (potentially slower) time course or is even absent. Moreover, caffeine-induced REM sleep differences were statistically not detectable after withdrawal over 44.5 h, indicating that a potential caffeine-related change in REM sleep promotion seems to be partly reversible. As withdrawal symptoms commonly reach peak intensity between 20 and 51 h after acute caffeine cessation (Juliano and Griffiths, 2004), volunteers have presumably already overcome the acute phase of caffeine withdrawal at the time of sleep initiation.

Caffeine-induced reductions of REM sleep propensity have earlier been demonstrated after administration of caffeine right before daytime sleep (Carrier et al., 2007; Carrier et al., 2009). However, the same outcome observed from 13.5 hours after the last intake in the present study suggest that caffeine concentration is not the only determinant to induce REM sleep changes. This notion receives further support as high dosages do not consistently evolve an effect on REM sleep [(Karacan et al., 1976; Nicholson and Stone, 1980; Robillard et al., 2015) vs (Bonnet and Arand, 1992; Drake et al., 2013)]. On the other hand, in line with the present data, caffeine-induced reductions of REM sleep features have consistently been observed when sleep was scheduled in the morning hours or at daytime (Carrier et al., 2007; Carrier et al., 2009). Thus, caffeine effects on REM sleep seem to depend on circadian factors. The importance of the circadian phase (i.e., during strong circadian REM promotion in the early morning) might be comparable to the impact on the homeostatic component of sleep-wake regulation, which seems to be particularly sensitive to the effects of caffeine under conditions of high sleep pressure (Roehrs and Roth, 2008; Snel and Lorist, 2011). Taken together, an effect of caffeine on sleep appears to be more easily detectable if the experimental design allows the sleep feature, i.e. REM sleep, to reach a certain level.

Changes in sleep pressure have been shown to interact with the circadian expression of REM sleep (Dijk and Czeisler, 1995; Wyatt et al., 1999) and might thus have modulated the present effects as the sleep episode was scheduled 5 h after volunteer’s habitual bedtime. However, it is important to note that we scheduled a one-hour nap episode in the evening prior to the sleep episode. In this nap participants slept for around 30 min, and sleep pressure as indexed by EEG SWA did not significantly differ between conditions (Weibel et al., 2020b). Moreover, additional analyses revealed no difference in all-night SWA and SWA of the first sleep cycle during nocturnal sleep after 16 h of wakefulness and the present reported sleep episode after 21 h of wakefulness among the three conditions (statistics not presented). Thus, our results suggest that the potential occurrence of increased sleep pressure did not strongly affect the present findings which can thus likely be traced back to the circadian component of sleep regulation.

Interestingly, volunteers in the caffeine condition indicated worse sleep quality indexed as more difficulties at awakening following sleep and feeling more tired compared to placebo. While SWS is commonly related with recuperation and subjectively perceived good sleep quality (Keklund and Akerstedt, 1997), the duration of REM sleep has been recently reported to be an important determinant for the perception of sleep quality (Della Monica et al., 2018). Thus, it is tempting to associate the changes of REM sleep with the perception of poorer sleep quality, although REM sleep did not account for the perceived sleep quality in the present sample. Moreover, the changes in sleep quality which were particularly evident at awakening in the present sample might also influence sleep inertia. Thus, the observed perception of non-restorative sleep might promote daily caffeine consumption in order to compensate the lack of feeling refreshed upon wake-up and thus act as a negative reinforcer.

Our results need to be interpreted taking the following limitations into careful consideration. First, based on the potential influence of the menstrual cycle on sleep-wake regulation (Shechter and Boivin, 2010) and the use of oral contraceptives on caffeine elimination (Abernethy and Todd, 1985; Balogh et al., 1995), female individuals were excluded which limits the generalizability of the present findings. Second, the effects of caffeine on sleep vary with age (Drapeau et al., 2006; Carrier et al., 2009; Robillard et al., 2015) and thus the present results only refer to young healthy individuals. Third, previous studies reported that a specific variation in adenosine receptor gene modulates the effects of caffeine on sleep (Rétey et al., 2007) which was however not controlled in the present study. Fourth, while we investigated REM sleep propensity at one circadian phase only, the optimal design would involve several circadian phases and a control for sleep pressure to allow conclusions on circadian regulation of sleep (such as in so-called forced desynchrony studies, see (Wyatt et al., 2004)).

In summary, we have first evidence that regular caffeine intake during daytime weakens REM sleep promotion, as evident in prolonged REM latency and reduced REM sleep accumulation. In the light of the evidence at present, these caffeine-induced changes in the circadian axis of sleep promotion may specifically occur when sleep is delayed and centered around the morning hours such as at the weekend or during shiftwork. Caffeine-induced REM sleep changes may contribute to a worse quality of awakening and thus reinforce the maintenance of daytime caffeine intake.

## Acknowledgements

We thank our interns Andrea Schumacher, Laura Tincknell, Sven Leach, and the study helpers for their help in data acquisition and all the volunteers for participating in the present study. We are grateful for the help in study organization by Dr. Ruta Lasauskaite and the medical screenings by Dr. med. Corrado Garbazza and Dr. med. Helen Slawik.

## Funding

The present work was funded by the Swiss National Science Foundation (320030_163058), the Matthieu Stiftung, the Nikolaus und Bertha-Burckhardt-Bürgin Stiftung, and the Janggen-Pöhn-Siftung.

## Declaration of conflicting interests

The authors declare that there is no conflict of interest.

## Notes

### Competing Interest Statement

The authors have declared no competing interest.

